# Quantifying influences on intragenomic mutation rate

**DOI:** 10.1101/2020.01.29.925719

**Authors:** Helmut Simon, Gavin Huttley

**Affiliations:** Research School of Biology, the Australian National University

**Keywords:** variance in mutation rate, context dependent mutation, recombination dependent mutation, ARMA models

## Abstract

We report work to quantify the impact on the probability of human genome polymorphism both of recombination and of sequence context at different scales. We use population-based analyses of data on human genetic variants obtained from the public Ensembl database. For recombination, we calculate the variance due to recombination and the probability that a recombination event causes a mutation. We employ novel statistical procedures to take account of the spatial auto-correlation of recombination and mutation rates along the genome. Our results support the view that genomic diversity in recombination hotspots arises from a direct effect of recombination on mutation rather than from the effect of selective sweeps. We also use the statistic of variance due to context to compare the effect on the probability of polymorphism of contexts of various sizes. We find that when the 12 point mutations are considered separately, variance due to context increases significantly as we move from 3-mer to 5-mer and from 5-mer to 7-mer contexts. However, when all mutations are considered in aggregate, these differences are outweighed by the effect of interaction between the central base and its immediate neighbors. This interaction is itself dominated by the transition mutations, including, but not limited to, the CpG effect. We also demonstrate strand-asymmetry of contextual influence in intronic regions, which is hypothesized to be a result of transcription coupled DNA repair. We consider the extent to which the measures we have used can be used to meaningfully compare the relative magnitudes of the impact of recombination and context on mutation.

---

Germline mutations are estimated to occur in humans with an average probability of 1.28 × 10^−8^ per site per generation, with ∼93% of these being point mutations (Jónsson *et al.* 2017; Roach *et al.* 2010). Germline point mutations result in the creation of single nucleotide variants (SNVs) in a population. Evidence of genomic heterogeneity in mutation has been predominantly derived from between or within species analysis of genetic variation. For instance, mutation heterogeneity is implicitly supported by genomic heterogeneity in substitution rates (Hodgkinson *et al.* 2009; Ying *et al.* 2010) and in the abundance of nucleotides (Cuny *et al.* 1981). Only recently have explicit *de novo* mutation studies (e.g. Smith *et al.* 2018) been reported, and these too support a heterogeneity in mutation processes. The mechanistic origins of this mutation heterogeneity remain unclear. Likely candidates include a direct mutagenic influence of meiotic recombination and the effect of sequence neighborhood. Analyses of these potential contributors have predominantly drawn on SNV analyses and have led to inconsistent conclusions. Here we focus on development and application of a consistent analytical framework to quantify the relative importance of these different factors.

It has been established that the rate of mutation is non-uniform along the genome of humans and other species. The phenomenon of mutation heterogeneity was first observed in the bacteriophage T4 prior to the availability of DNA sequencing (Benzer 1961). Subsequent DNA sequence analyses of homologous genes revealed that G and C nucleotides were far more mutable than A and T nucleotides (Gojobori *et al.* 1982) and that mutation rates at these sites are influenced by neighbouring bases (Bulmer 1986). We now have evidence that such non-uniformity can occur at scales ranging from individual nucleotides to multi-megabase sized regions (Hodgkinson and Eyre-Walker 2011). For instance, the heterogeneity of DNA composition suggests the existence of mutation rate heterogeneity at megabase scales and this has been supported by *de novo* mutation studies (Smith *et al.* 2018). A more complete quantitative understanding of the various mechanisms that may influence mutation rate and hence lead to mutation rate variability remains the subject of active research activity.

Previous studies have identified a number of key determinants of mutation rate, prominent among which are recombination rate and sequence neighbours. However, these studies have differed in certain of their conclusions. For example, relatively recent studies of *de novo* mutations have provided strong evidence of a direct causative effect of recombination on mutation (Arbeithuber *et al.* 2015). That nucleotide diversity is higher in regions of high recombination has been known for some time (Lercher and Hurst 2002; Duret and Arndt 2008). Whether this reflects a direct effect of recombination on mutation or an influence of selective sweeps in reducing diversity in regions of lower recombination is disputed (e.g. Kern and Hahn 2018). Previous population analyses applied to this problem used linear regression models (Lercher and Hurst 2002; Duret and Arndt 2008; Mugal and Ellegren 2011). Estimates from these approaches are potentially problematic as the methods used do not control for spatial auto-correlation of recombination and mutation rates across the genome.

The hypermutability of CpG dinucleotides (and the preponderance of genetic variation within this context) exemplifies the important influence of sequence context on the rate of mutation. In mammals and some other species, the transition mutation C→T, where the C is part of a CpG dinucleotide, is several times more common than mutations at other sites (Ehrlich and Wang 1981). The biochemical cause is known to be the spontaneous deamination of the highly unstable methylated cytosine base 5-methylcytosine (Coulondre *et al.* 1978). In mammals, methylation of C is highly context dependent, occurring almost exclusively at CpG dinucleotides (Ramsahoye *et al.* 2000).

It has been demonstrated that all point mutations are affected to a greater or lesser effect by sequence context (Zhu *et al.* 2017). Using a log-linear model, Zhu *et al.* (2017) dissected the influence of nucleotide distance and the joint versus independent influence of multiple nucleotides. These authors argued that the dominant neighbourhood influences lay within ±2 for transition mutations, ±3 for transversions. Zhu *et al.* (2017) did not, however, directly address mutation rate or variance in the sense described above. An analysis using the *R*^2^ metric of a linear model to measure the contribution of different contexts to variance (Aggarwala and Voight 2016) argued that nucleotides up to 3 sites distal can have a major influence on mutation rates. The linear regression model used by Aggarwala and Voight is inappropriate in the sense that it does not yield the maximum likelihood estimates of model parameters for this data, due to the binomial nature of the sampled data and the condition of heteroscedasticity consequently not being satisfied (Agresti 2002, p. 120). Previous approaches also did not address issues of bias arising from neighborhood size. Bias will tend to inflate estimates of variance as a given data set of mutation counts is further subdivided into “buckets”, the number of which increases with neighborhood size *k* at the rate 4^*k*^.

One approach to quantifying the relative contributions of different factors on mutation is to measure the proportion of variance in mutation rate explained by them. Conversely, this measurement also indicates how much variance remains unexplained. Inherent in discussion of the variability of mutation rate is the assumption that each site in the genome has a specific mutation rate. Hence, we define the “total” variance in mutation rate as the conventional statistical variance of these quantities. This variance has been estimated by comparing variable positions in orthologous alignments of closely related species such as humans and chimpanzees (Hodgkinson *et al.* 2009). The probability of an SNV at a site is assumed to be some multiple *r* of the site mutation rate, with *r* fixed in each population. (The underlying mutation rates are assumed to be the same in humans and chimpanzees.) The variance in mutation rate can then be calculated from the number of SNVs that are observed at orthologous sites in both sequences. The conclusion from this approach was that there was substantial variance in the human mutation rate (∼64% of total variance) that was not explained by the interaction of a base with its immediately adjacent nucleotides (Hodgkinson *et al.* 2009). These authors largely dismiss the potential role of larger sequence contexts, a conclusion that is challenged by the results of other studies (Zhu *et al.* 2017; Aggarwala and Voight 2016).

Here we report work quantifying the contribution to the probability of human genome polymorphism that can be attributed to recombination and to sequence context at different scales. We use a Bayesian approach to quantify the uncertainty in our estimates of the variance and to overcome issues of bias which occur if a conventional estimator were used instead. Our results produce estimates of recombination induced mutation that are consistent with those from *de novo* mutation studies. We further establish that when considered across all point mutations, the influence of sequence neighbourhood is dominated by 5-mer effects reflecting the markedly greater relative abundance of transition mutations. Finally, we emphasize the complexity in comparing the contributions to mutation of a state (sequence context) versus the contribution to mutation of an event (recombination). Overall, we establish that a substantial proportion of mutation heterogeneity remains unaccounted for.

## Materials and methods

### Data

Data on variants was sampled from Ensembl release 89 (for influence of context) and release 92 (for influence of recombination) variation databases (Cunningham *et al.* 2015) using the query capabilities of ensembldb3 (Huttley and Ying 2009). Human variants were restricted to those identified by the 1000 Genomes (1KG) Project (Aut 2015), but without restriction by source population.

deCODE provides estimated recombination rates averaged over 10-kilobase (kb) blocks. The files female_noncarrier.rmap, male_noncarrier.rmap and sex-averaged_noncarrier.rmap were downloaded from https://www.decode.com/addendum/ (Kong *et al.* 2010). These correspond to female, male and sex-averaged standardized recombination rates respectively. The hg18 genome coordinates were mapped to GRCh38 using pyliftover, a Python implementation of UCSC LiftOver (Tretyakov 2013).

### Variance in probability of polymorphism due to recombination

In order to estimate variance in the probability of mutation that can be explained by recombination or by sequence neighbourhood, we employ the SNV density as a surrogate. Counts of SNVs within the 10-kb blocks defined by deCODE were determined from the Ensembl variation database records. We excluded blocks where no SNVs were reported in Ensembl and blocks that were identified by deCODE as overlapping unsequenced regions.

The relationship between recombination rate and SNV density may be confounded by spatial auto-correlation of these quantities along the genome. The impact of auto-correlation on the residuals of a linear model was confirmed by plotting the covariances of the residuals for blocks separated by up to 50 blocks using statsmodels (Seabold and Perktold 2010) (Figure S1). Allowing for auto-correlation in our model requires maintaining the lags between the 10-kb blocks and thus it was necessary to adjust regions with missing data. This was done using the Last Observation Carried Forward (LOCF) method (Molenberghs *et al.* 2014, p. 38). That is, for successive blocks excluded by our missing data criteria, SNV and recombination data from the immediate 5’ neighbour block were repeated.

Selection of the appropriate time-series model for the residuals depends on whether their distribution is stationary. The statsmodels (Seabold and Perktold 2010) implementation of the augmented Dickey-Fuller test (Mills 2008, p. 79) was used to demonstrate stationarity of the residuals. (See also Figure S2.) Stationarity allows us to apply Wold’s decomposition theorem (Mills 2008, p. 12) to conclude that the residuals can be approximated by an auto-regressive moving average (ARMA) model of some order (*p, q*) where *p* and *q* are non-negative integers and *p* > 0. Optimal values of *p* and *q* were chosen by evaluating models for *p* ≤ 10 and *q* ≤ 4 using the statsmodels (Seabold and Perktold 2010) ARMA implementation to find which had the lowest value of the Aikake Information Criterion (AIC). (In the case of chromosome 9, the model with the second lowest AIC was used, as the lowest model confounded the subsequent Markov Chain Monte Carlo step.)

A Bayesian Markov Chain Monte Carlo (MCMC) approach implemented in the software package PyMC3 (Salvatier *et al.* 2016) was used to simultaneously estimate the slope, intercept and *p* + *q* ARMA parameters. This was developed to provide a more robust approach than iterative adjustment of the parameters (Mizon 1995) as is undertaken with, for example, the Cochrane-Orcutt procedure. The intercept *α* obtained from this process represents the model’s prediction of SNV density for genomic segments with a recombination rate of zero. Therefore, given the average SNV density 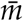, we can estimate the proportion of SNVs, and hence of mutations, caused by recombination as 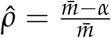. Then if *x* is the number of mutations that occur in 1 Mb of DNA sequence in a specific generation and *y* is the number of recombination events occurring in that sequence in the same generation, 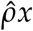 is the expected number of new mutations in that segment caused by recombination. Since they must be caused by recombination events occurring in that generation, the expected number of mutation events per recombination event is 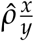. Therefore multiplying 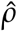 by the ratio of mutations per generation to recombination events per generation gives the average number of mutations produced by each recombination event. The estimated variance in SNV density due to recombination 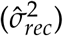 is calculated as the difference between the total variance in SNV density and the sum of squares of the residuals. The ratio of this quantity to total variance in SNV rate is the proportion of variance in SNV rate attributable to recombination (*R*^2^).

Our model does not take account of error in the estimation of recombination rates in the blocks. To determine the impact of this, we tried adding a normal perturbation of the recombination rates to the model. This made little difference to the posterior distribution, which we hypothesize is due to averaging the recombination rates over a large number of blocks.

### The variance in SNV density conditioned on context

We estimated the probability of polymorphism for all point mutation directions from all sequence contexts of size *k* that contained a central point mutation. The mean and variance can be obtained from these in a straightforward manner. The variance conditioned on different central bases or different point mutation directions can be measured by filtering the appropriate subset of the data. We now expand on our model and approach to estimation of the variance.

We denote by 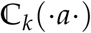 the set of 4^*k*−1^ sequence contexts with central base *a* ∈ {*A, C, G, T*}. The union, 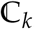, of the four such sets contains the 4^*k*^ distinct *k*-mer sequences. As we are concerned with neighbourhoods centered on a mutating base, *k* is an odd numbered integer with values of 3, 5, 7 or above.

For a sequence *S*, our model assigns to each site a fixed probability *m* of being polymorphic for an SNV and assumes that the mutation events for different sites occur independently (see Assumptions below). It is the variability of *m* that can be explained by context that is the object of the analysis.

For a context *c*, let *p*_*c*_ be the proportion of sites in *S* matching it. We denote by *m*_*c*_ the probability that a randomly selected site matching the context *c* will have a SNV at the central base. Then *m*_*c*_ is the average SNV probability over the sites with context *c*. We denote by 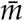 the average SNV probability over the entire sequence. For any *k* we have:

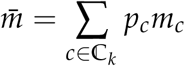

Then the total variance in SNV density accounted for by sequence neighbourhoods of size *k* is:

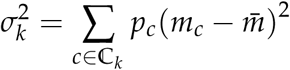

This total variance can be partitioned into components consisting of variance attributable to each point mutation *a* → *b* as

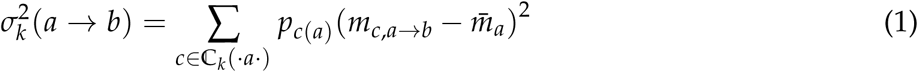

where *p*_*c*(*a*)_ is the proportion of sites with base *a* whose context matches *c*; *m*_*c,a*→*b*_ is the probability of polymorphism arising from mutation of base *a* to base *b* in context *c*; and 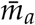 is the probability that a site with base *a* will have an SNV.

We consider the proportion of contexts *p*_*c*_ (and *p*_*c*(*a*)_) as a fixed or known quantity, as contexts can be counted exactly with reasonable efficiency. We then estimate the values *m*_*c*_ by 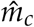, the empirical SNV density in context *c*. The estimated value of 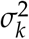 is then given by:

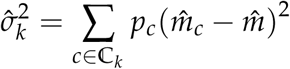

where 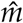 is the empirical SNV density for the entire sequence. A similar equation applies if we condition on a specific point mutation direction *a* → *b*. For instance, we can further condition on C sites with 5′ G and G sites with 3′ C in order to isolate the CpG effect.

### Assumptions

Some of the assumptions made in the above model may be invalid in practice. We deal with this by filtering these conflicting cases from the data, as follows.

We have assumed that each site has a fixed probability of being polymorphic and that the resultant Bernoulli distributions are independent between sites. These assumptions fail if a site mutates more than once, since we allow the nucleotide to influence mutation rate. It similarly fails if a neighbouring site mutates, since we allow context to influence mutation. We therefore only include those SNVs in our data set which are biallelic, with one allele being the ancestral allele; and for which there are no variants in the immediate neighbourhood (4 bp on either side). (It is recognized that this does not eliminate the case in which subsequent mutations have occurred within the context and achieved fixation).

### Bayesian model for estimation of variance due to context

We use Bayesian conjugate priors to derive a posterior distribution for each instance of mutation direction within a particular context (e.g. ACT → ATT mutation in the case of 3-mers). For each such case we have a count of *k*-mers (number of “trials”) and a count of variants at the central base of the *k*-mer (number of “successes”). The probability of polymorphism is given by estimating the probability parameter of a binomial distribution on these quantities. The conjugate prior for the binomial distribution is the beta distribution, so we use a Beta(1,1) distribution as a prior. We thus derive a posterior beta distribution for the mutation rate for the cell. We generate samples of the posterior distribution for the variance due to context by generating samples for the probability of polymorphism for each cell from the beta distributions and applying the right hand side of equation (1) to the samples to generate samples for the weighted variance.

This method requires that the number of mutation type and context pairs having no variants in the data is small. For such cells the posterior distribution on mutation rate would be Beta(1,1), the uniform distribution on [0, 1] and hence the variance in mutation rates would be inflated.

## Data and Software Availability

All raw and processed data used in this work are available at https://zenodo.org/record/3608379. Scripts and Jupyter notebooks developed specifically to perform the data sampling and analyses reported in this work were written in Python version ≥3.5 and are freely available under the GPL at https://github.com/helmutsimon/ProbPolymorphism and at https://zenodo.org/record/3628923.

Dependencies included cogent3 (Knight *et al.* 2007), ensembldb3 3.0a1 (Huttley and Ying 2009), pymc3 3.6 (Salvatier *et al.* 2016), theano 1.0.4 (Theano Development Team 2016), pyliftover 0.3 (Tretyakov 2013), sqlalchemy 1.2.2 (Bayer 2012), scipy 1.2.1 (Virtanen *et al.* 2019), numpy 1.15.1/ 1.16.3 (Virtanen *et al.* 2019), pandas 0.23.4/ 0.24.2 (McKinney 2010), statsmodels 0.10.1 (Seabold and Perktold 2010), scikit-learn 0.21.2 (Pedregosa *et al.* 2011), click 6.7 (Ronacher 2009), scitrack 0.1.3 (Huttley 2016), matplotlib 3.0.3 (Hunter 2007) and seaborn 0.9.0 (Waskom *et al.* 2017).

## Results

We estimated the contributions of recombination and context to the variance in SNV density using data from the Ensembl variation database (Cunningham *et al.* 2015). The SNV density for a sequence is defined as the number of qualified SNVs in the sequence divided by the sequence length. Only 1KG Project (Aut 2015) variants were considered in the interests of consistency in SNV discovery. The point mutation direction from which a SNV was derived was inferred using the ancestral nucleotide state as recorded in Ensembl.

### Effect of recombination on SNV density

We evaluated the relationship between recombination and mutation rates using linear regression. Our aim was to recover the slope and intercept parameters from which other quantities of interest can be inferred. The slope parameter gives us the increase in SNV density for a given increase in recombination rate. In particular, a positive slope parameter indicates a positive effect of recombination on SNV density and hence mutation. The intercept parameter is the value of SNV density corresponding to a recombination rate of zero under the model. The estimated variance in SNV density due to recombination, which we denote by 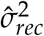, is calculated as the difference between the total variance in SNV density and the sum of squares of the residuals. The ratio of this quantity to total variance in SNV rate is the proportion of variance in SNV rate attributable to recombination. This ratio is the standard metric *R*^2^ (coefficient of determination), which measures the fit of a linear model in terms of explained variance in the observed data.

In modelling the influence of recombination, we assumed that each recombination event has some probability of giving rise to a mutation. We used a partitioning of the genome into 10-kb segments for which average sex-averaged recombination rates were available (Kong *et al.* 2010). These rates are normalized relative to the average genetic distance over all of the 10-kb bins of 0.0116 centimorgans. SNV densities were derived from the number of SNVs in a segment. This quantity is expected to vary linearly with the probability of a recombination event occurring in the segment (that is, with the genetic length of the segment).

We began by fitting an ordinary least squares linear regression (OLSLR) model to the data. Use of an OLSLR model for inference requires residuals to be mutually independent, in particular that there is no correlation between adjacent bins along the genome (spatial auto-correlation). By analysing the residuals from an OLSLR model we identified a high level of auto-correlation (see Supplementary Figure S1) and determined that they were most appropriately modelled by an ARMA(*p, q*) model, where *p* and *q* are non-negative integers and *p* > 0 (see Materials and methods). For each chromosome, we tested a range of ARMA models to find the one with the lowest Aikake Information Criterion (AIC) score for the data. The slope, intercept and ARMA error parameters were simultaneously estimated using a Bayesian Markov Chain Monte Carlo (MCMC) approach (see Materials and methods), obviating the need for iterative “adjustment” steps.

The above process was applied to all odd-numbered chromosomes individually. The estimates for variance in SNV rate due to recombination 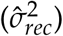 are shown as violin plots in Figure 1. It can be seen that there were some significant differences in the variance estimates for different chromosomes. In particular, chromosomes 9, 15 and 17 show significantly higher levels of variance in SNV density due to recombination. There were also significant differences in estimates of the slope and intercept parameters (see Table S1). These differences between chromosomes precluded estimation of the influence of recombination across the genome as a whole. Specifically, using a model which set the slope and intercept parameters to be common across chromosomes while allowing differing ARMA models and parameters for each chromosome resulted in a y-intercept that was larger than the average SNV density, which is inconsistent with results from individual chromosomes. Modifications of this approach that used the sex-specific recombination maps did not result in any substantial differences (results not shown).

**Figure 1.**
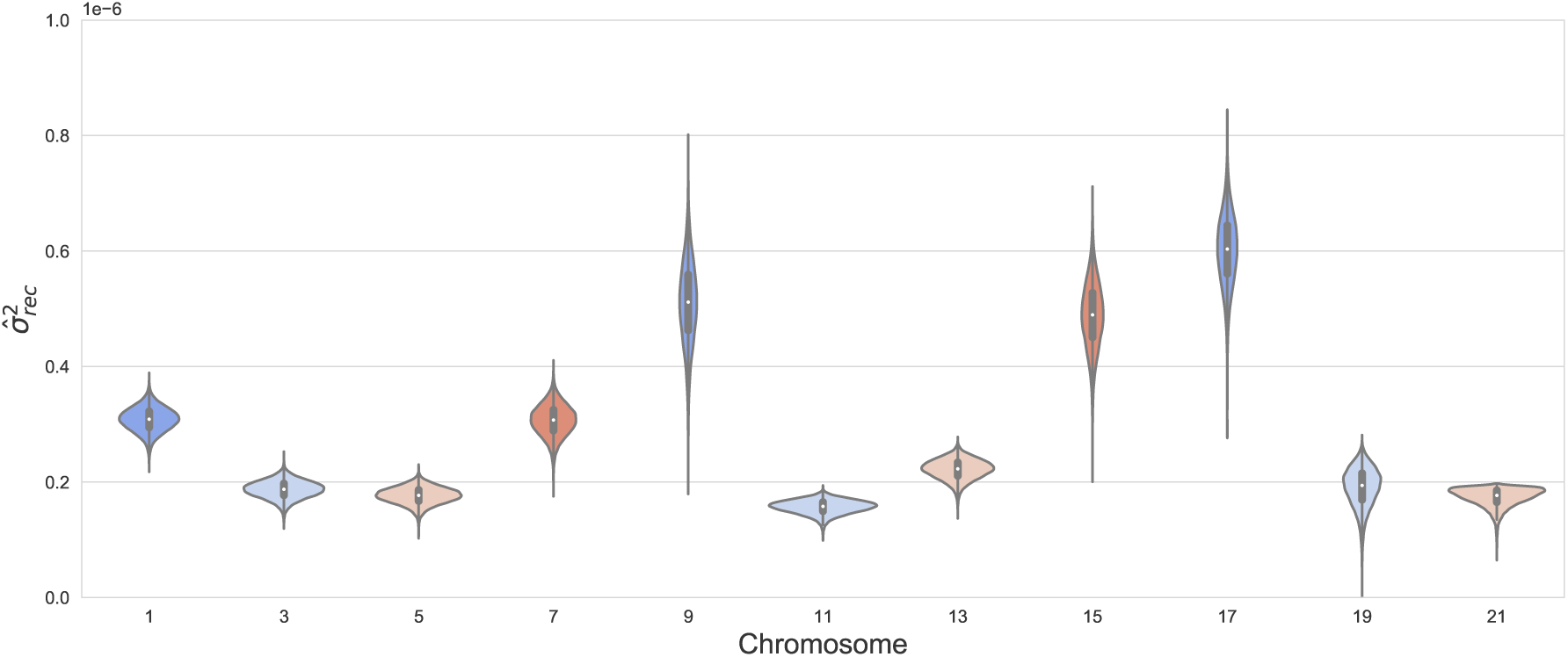
Estimated variance in SNV density attributable to recombination by chromosome. The variances 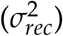 are estimated by fitting a linear model to each chromosome, with residuals modelled by an ARMA(p,q) model optimised for each chromosome. The variance due to recombination is the difference between total variance and the variance not explained by the model.

Estimates for the slope (change in SNV density per centimorgan) provided strong evidence for a positive effect of recombination rate on SNV density across all chromosomes. The estimates ranged from 0.0067 for chromosome 21 to 0.0091 for chromosome 17 (see Table S1). The corresponding 95% credibility intervals (hereafter CI) of these estimates were 0.0042-0.0091 and 0.0068-0.011 respectively. For all chromosomes tested, the posterior probabilities that the slope was ≤ 0 were ≤ 10^−4^ (estimated from the MCMC variates).

In the linear model, the y-intercept represents the predicted SNV density for a recombination rate of zero. Estimates for the y-intercept ranged from 0.0251 (95% CI of 0.0247-0.0255) for chromosome 1 to 0.0274 (95% CI of 0.0271-0.0277) for chromosome 21. The difference between the mean SNV density and the y-intercept parameter is more significant as it represents the difference between the average observed SNV density calculated and the observed data and the SNV density predicted for a recombination rate of 0. That is, this difference measures the part of the SNV density due to recombination. Dividing the difference by mean SNV density gives the proportion of SNVs due to recombination (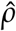, Materials and methods). This quantity varied between 0.0032 for chromosome 5 and 0.0051 for chromosome 17. These chromosomes are also at the extremes of variance due to recombination.

We also examined the extent to which the effect of recombination on SNV density differed for the 12 point mutations. We carried this analysis out separately for all odd numbered chromosomes. As an example, results for chromosome 1 are shown in Table 1. We accepted that recombination has had a positive effect on mutation when the posterior probability that the slope was less than zero was found to be less than 0.05. On this basis, the mutations for which recombination influenced mutation in Chromosome 1 comprise all four transitions (C→T, T→C, A→G, G→A) and the N→S transversions C→G, G→C, T→G and A→C.

**Table 1.**
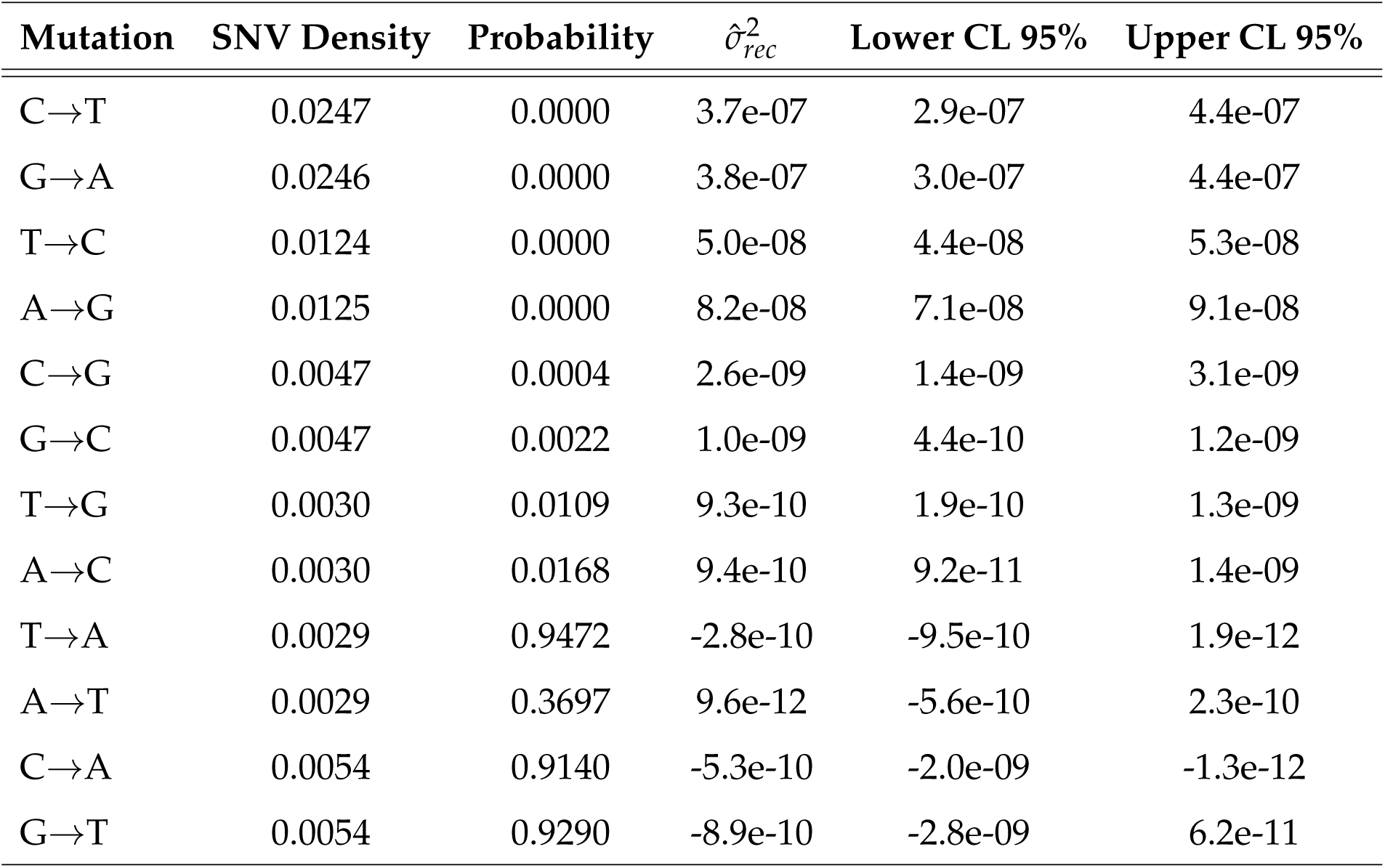
Analysis of the linear relationship between recombination rates and SNV densities for chromosome 1 disaggregated by mutation direction. ‘SNV Density’ is the SNV density for that mutation direction (conditioned on ancestral allele); ‘Probability’ is the posterior probability that the slope parameter from the linear regression is less than zero; 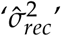 is the estimated variance due to recombination and ‘Lower CL 95%’ and ‘Upper CL 95%’ are the limits of the 95% credibility interval for 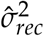. Since the estimated variance in SNV density due to recombination is calculated as the difference between the total variance in SNV density and the sum of squares of the residuals, it will be negative if the model fit is worse than for a line with zero slope. This is likely to occur when the ‘Probability’ value is significantly greater than zero and we reject the model.

For SNVs derived from transition mutations, evidence for an association with recombination rate was consistent across all chromosomes (Figure 2 and Supplementary Table S2). This was not the case for the transversion mutations. For SNVs derived from transversions, evidence of an influence of recombination ranged from inconsistent to none. For instance, for transversions to G/C, the posterior probabilities for most chromosomes met our 0.05 threshold. In contrast, there was no evidence of an influence of recombination on transversions to A/T for most chromosomes. Additionally, if a mutation type appears to be influenced by recombination, so does its strand-symmetric counterpart. However, the values for variance due to recombination for a mutation and its strand-symmetric counterpart, while of similar magnitude, do not necessarily coincide, even using the 95%CI. For all chromosomes the mutations with the highest variance due to recombination are C→T and G→A, the same as are subject to the CpG effect.

**Figure 2.**
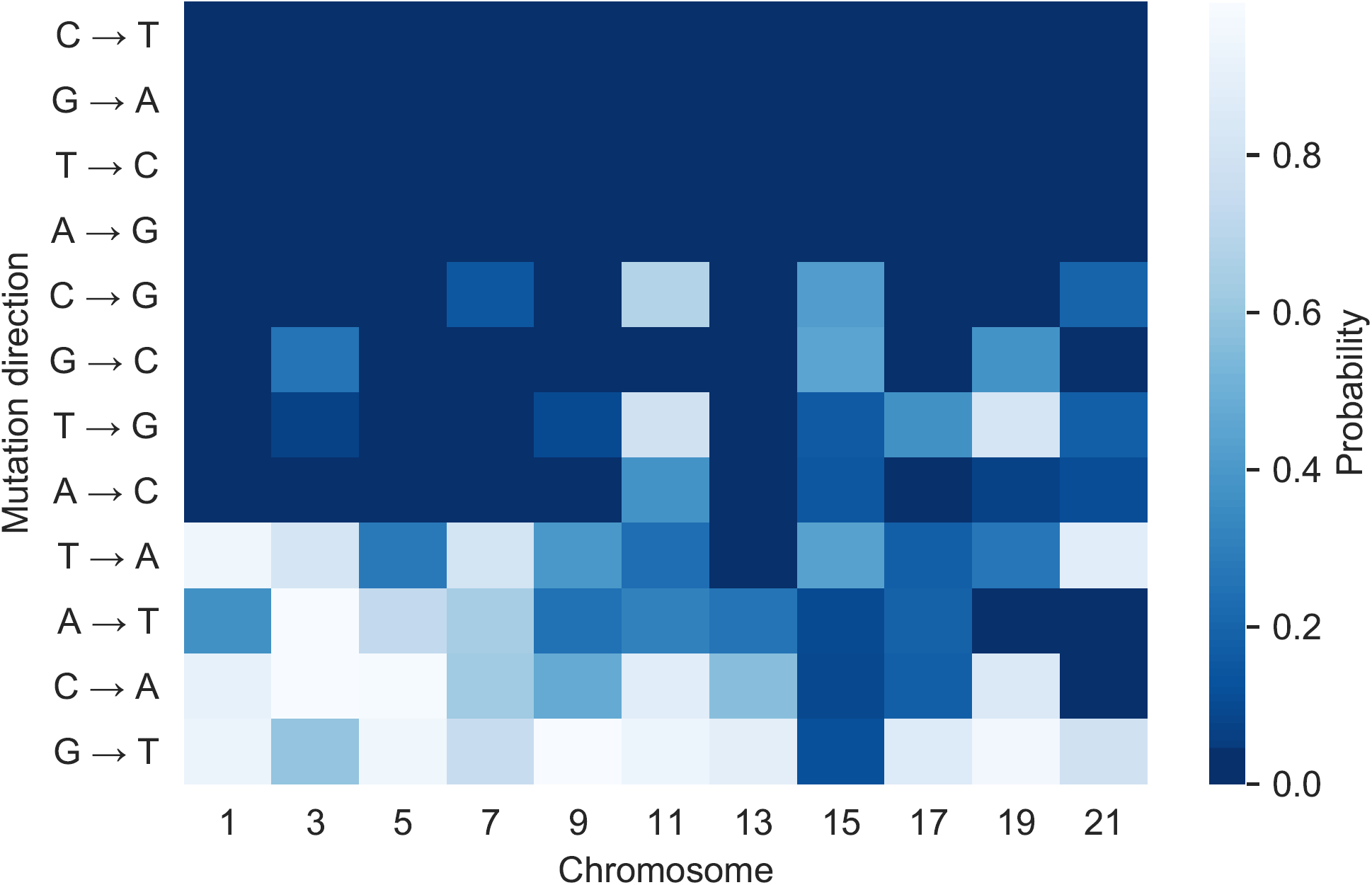
Evidence for the effect of recombination on mutation by mutation direction and chromosome. ‘Probability’ is the posterior probability that the slope parameter from the linear regression is less than zero. A darker shade indicates a high probability that SNV density has a positive linear relationship with SNV density. Cells meeting the 0.05 threshold are in the deepest blue.

### Variance in SNV density due to context

In our analysis of variance in SNV density due to context, we restricted ourselves to intronic 1KG Ensembl variants, to reduce confounding due to selection. All SNVs and contexts were oriented with respect to the annotated strand of the gene. We used data from all odd-numbered autosomes. For consistency with our assumption that each site has a fixed mutation rate, only biallelic variants separated by ≥ 4 nucleotides from another SNV were considered. Rather than use a conventional sample or plug-in estimator of the average SNV density for each context, we worked with samples from a posterior (beta) distribution to the binomial likelihood function. This allowed us to sample and graph posterior distributions for variance due to context, showing the uncertainty in the parameter estimates (see Materials and methods). It also allowed us to calculate the (posterior) probabilities of particular conditions by counting the proportion of MCMC variates satisfying the condition.

To evaluate the relationship between sequence context and the probability of a SNV requires further definition of SNV density. For a specific sequence context of size *k* (including the middle position), there are 4^*k*^ distinct contexts (hereafter *k*-mers). To illustrate calculation of SNV density, consider the 3-mer ACA. We estimated the SNV density for ACA as the number of occurrences of ACA for which the middle position had an SNV divided by the total number of occurrences of the *k*-mer ACA. This can be further partitioned into the different point mutations from C. The estimated variance attributable to sequence context of size *k*, which we denote by 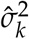, is thus the variance computed across all 4^*k*^ such densities (see Materials and methods).

The value of 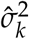 for *k* = 1,3, 5 and 7 is shown by the blue bars in Figure 3 for 1-mers to 7-mers. The case of *k* = 1 shows variance conditioned solely on ancestral base. We necessarily observe an increase in variance with increasing *k*, but the increments diminish markedly after 3. It is noteworthy that the variance due to the central base alone only comprises ∼ 11% of the variance due to 3-mers. The total variance due to 7-mers is ∼ 17% greater than that due to 3-mers. Variance in SNV density calculated in this way includes the influence of the central mutating base itself and the interaction between that central mutating base and its neighbourhood. To investigate the relative influence of these elements further, we show, using the brown bars, the values of 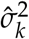 marginalised over the central base (Figure 3). This corresponds to evaluating the influence of the flanking nucleotides alone, by averaging out the effect of the central mutating base. We see that while these values are much lower than the unmarginalized values, the relative magnitude of the increments from 3-mer to 5-mer and from 5-mer to 7-mer are larger. We can conclude that the greater part of the unmarginalized variance due to 3-mers is explained not by the independent actions either of the central base or of the flanking bases, but by the interaction of the central base with its immediately adjacent neighbours. Furthermore, this interaction between a mutating base and its immediate neighbours is the largest contribution to variance for all values of *k* considered. As would be expected, a large component of this is due to the CpG effect (including its strand-symmetric counterpart) which we estimated as 0.00026, ∼ 60% of the variance due to 7-mers.

**Figure 3.**
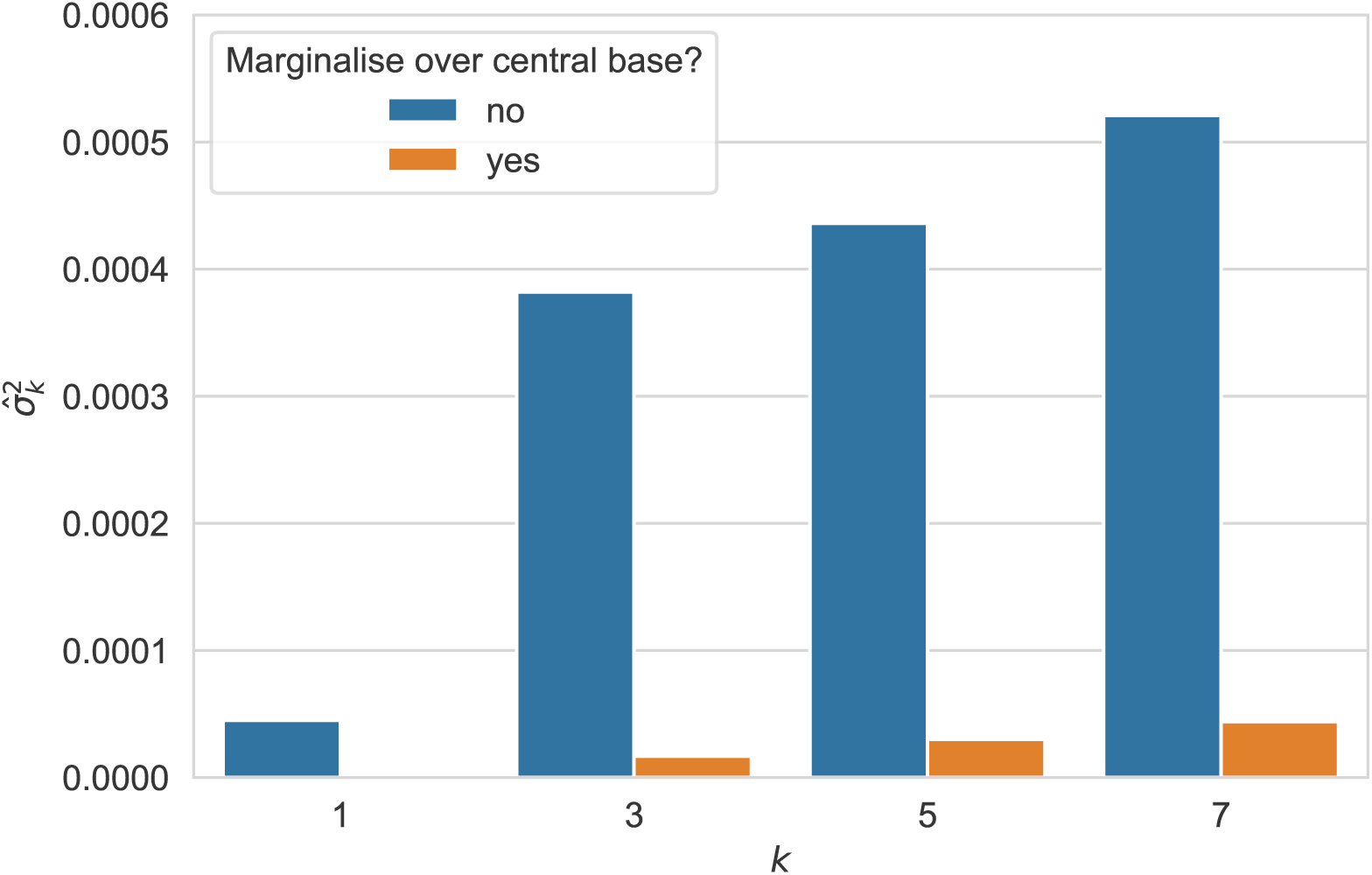
The interaction between a mutating base and its neighbourhood dominates variance in SNV density. The component of 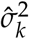 attributable to neighbourhood alone (i.e. marginalised over the central base) is shown in tan. This contrasts with the full value of 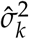 (shown in blue), which includes the interaction between a mutating base and its neighbourhood.

We also analysed the variance due to context for each of the point mutations separately. The results are shown in Figure 4 as posterior distributions for the variance due to context. (Values for the posterior mean are shown at Supplementary Table S3.) The increment in variance from 5-mer to 7-mer is greater than or approximately equal to that from 3-mer to 5-mer for all point mutations, with the exceptions of T→C / A→G transitions. In the case of transversions, the variance due to 7-mers is two or three times that due to 3-mers. The strong relative influence of 7-mers and 5-mers for transversions may appear to be at odds with the results aggregated over point mutations (Figure 3). However, since Figure 4 considers each point mutation separately, the central or ‘from’ base is fixed and the interaction between the central base and its immediate neighbours does not make a contribution. Thus the impact of increasing *k* is more similar to that of the marginalised quantities (Figure 3).

**Figure 4.**
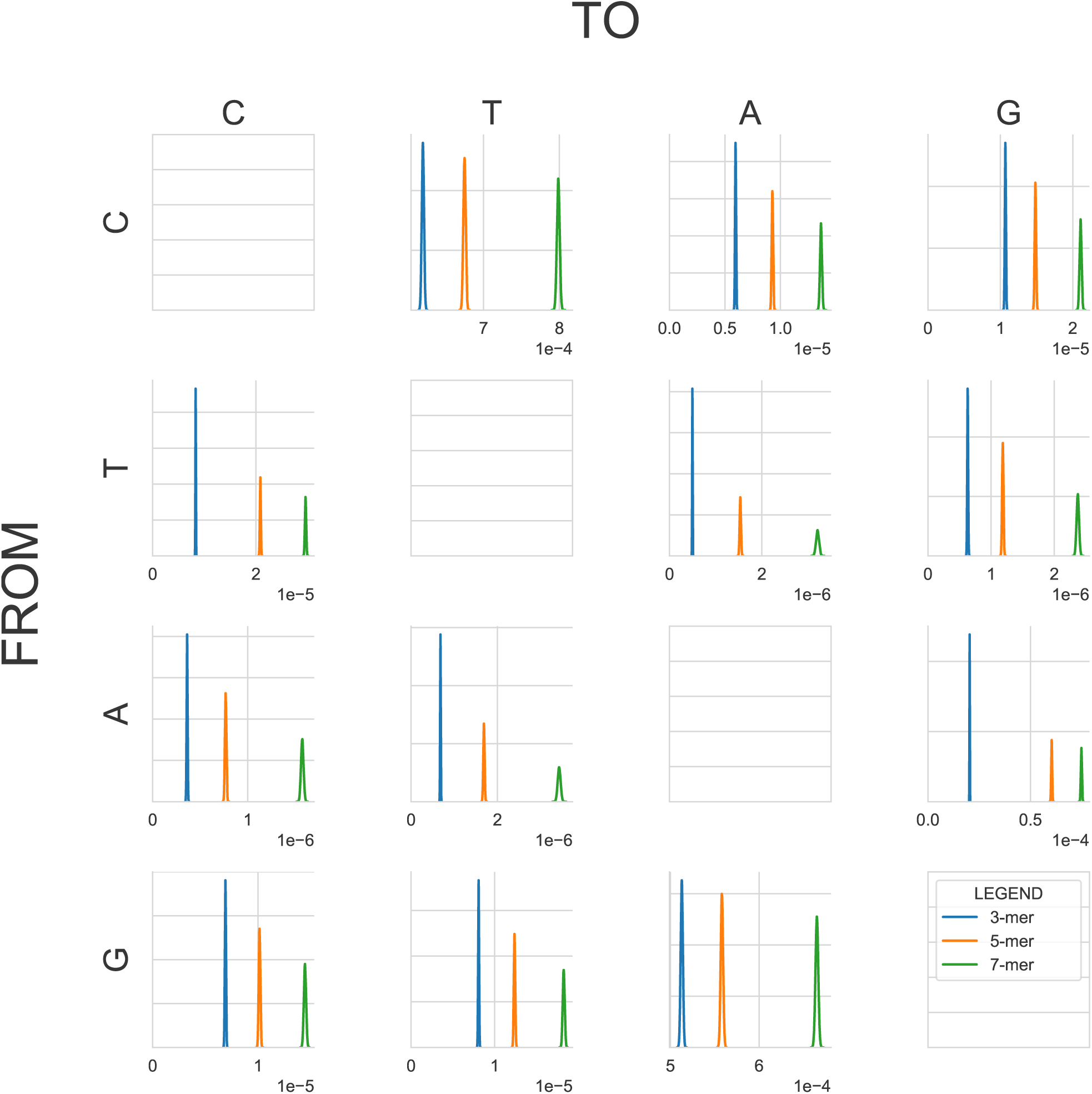
Posterior distributions of the variance of SNV density (x-axis) conditioned on 3-mer, 5-mer and 7-mer contexts for each of 12 mutation profiles. The Row/Column labels correspond to the *from* and *to* nucleotides. Note that the x-axes 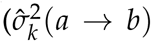, estimated variance due to context) and y-axes (probability density) scales vary between plots. In particular the x-axes for C→T and G→A mutations do not include the origin.

Examination of Figure 4 suggests that contextual influence does not always operate in a strand-symmetric manner. We investigated this further by plotting intronic mutations together with their strand-complements for the 7-mer case (Figure 5a). This demonstrates evidence of strand-asymmetry for all mutation types. This was especially marked for T→C / A→G transitions. Our criterion for rejecting strand-symmetry was that the 97.5 percentile of one of a pair of strand-complementary mutations was less than the 2.5 percentile of the other. As a control, we performed a similar analysis for intergenic regions. The results (Figure 5b) are generally consistent with the operation of strand-symmetric processes in intergenic regions. The pair of mutations G→T and C→A appear to be strand-asymmetric by our criterion and may be an exception or an artefact.

**Figure 5.**
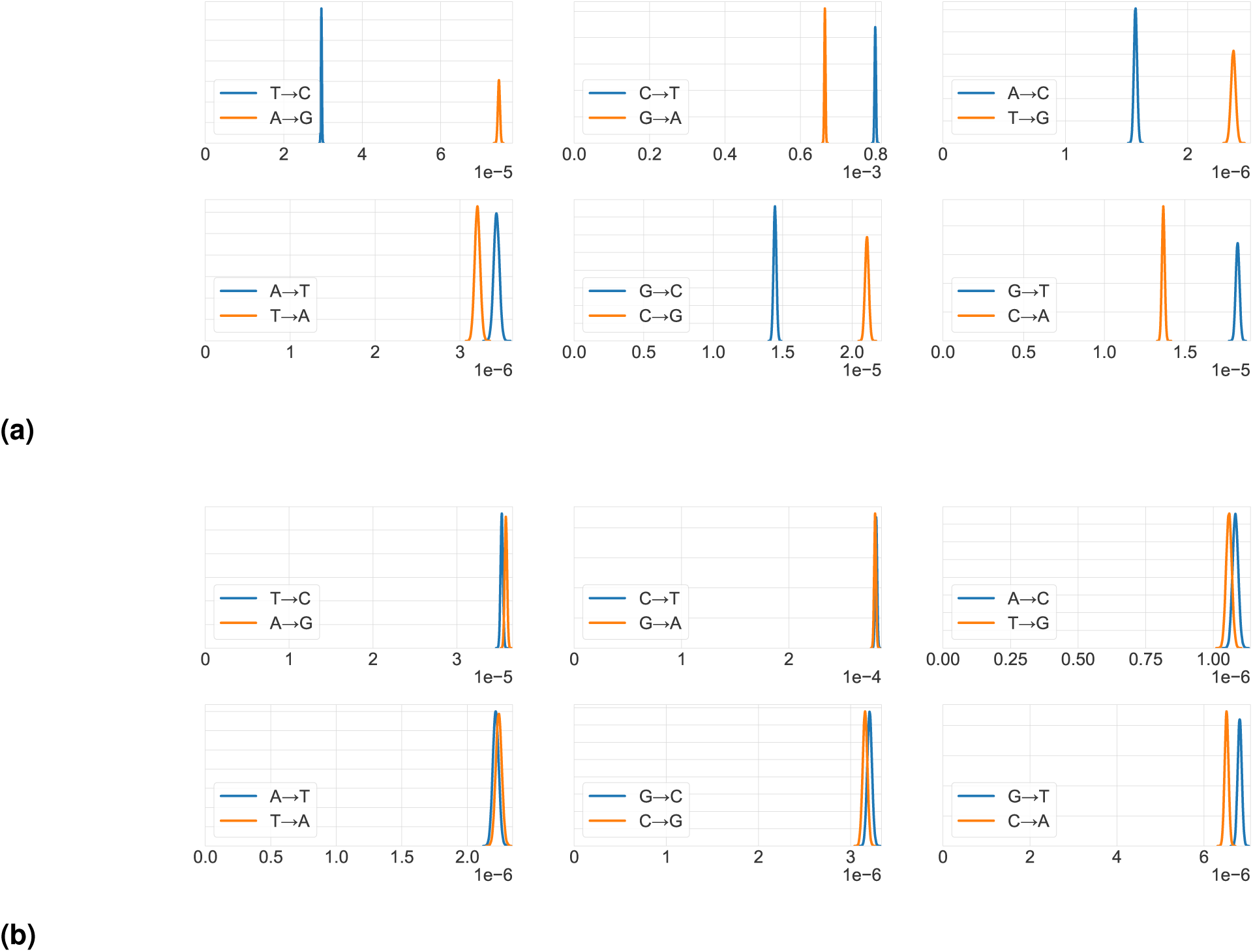
Variance in SNV density for specific mutation directions is strand-asymmetric for intronic regions, but strand-symmetric for intergenic regions. We show posterior distributions of 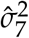 for each of the 12 mutation directions (x-axis), with strand-symmetric pairs shown on the same axes. The y-axes show probability density. (a) Intronic regions. (b) Intergenic regions.

## Discussion

A multitude of processes contribute to affect spontaneous mutagenesis. One approach to establishing the relative importance of these uses genomic heterogeneity in SNV density which is presumed to arise from the non-uniform action of factors contributing to mutation rate. Identifying an association between candidate factors and SNV density provides evidence of their effect on mutation. In this paper we developed an approach of estimating the variance in SNV density conditioned on either recombination rate or on sequence context of various sizes. We found an association between recombination and SNV density, obtaining estimates consistent with those from analyses of *de novo* mutation data and associated recombination events. Our analyses of contextual influences on SNV density demonstrated that the effect of context size differed between transition and transversion point mutations and did not always operate in a strand symmetric manner.

### The influence of recombination on mutation

A consistent positive association between recombination rate and SNV density was evident. We tested the hypothesis of a relationship between recombination and SNV density by testing for an association between the average recombination rate evident in 10-kb genomic segments with the SNV density in those segments. Due to substantial auto-correlation of mutation and recombination rates between the segments (Figure S1), we derived estimates of the variance using a linear regression model that was modified to incorporate this auto-correlation. For all chromosomes examined, the posterior probabilities that the slope was ≤ 0 were ≤ 10^−4^. This provides strong evidence of a positive association.

We determined the ratio between the estimated point mutation rate in the human genome (Jónsson *et al.* 2017) and the average SNV density obtained from our data. Applying this ratio enables us to express results in terms of mutation rates rather than SNV densities. On this basis, our estimates for the slope in terms of the base pair mutation rate per centimorgan ranged from 2.13 × 10^−9^ (chromosome 21) to 4.06 × 10^−9^ (chromosome 17) (Table S1).

A natural measure for the mutational effect of recombination is the probability that a recombination event causes a mutation event. Using values for average mutation rate (Jónsson *et al.* 2017) and recombination rate (Kong *et al.* 2010), we estimated that each recombination event results in an average of approximately 0.004 mutation events (see Materials and methods and Table S1). This estimate has a high degree of uncertainty, but was reasonably consistent across chromosomes, staying in the range 0.0031 - 0.0049 (except for chromosome 21 at 0.0022). The estimate is in good agreement with results obtained by a recent study of *de novo* mutations, which arrived at a figure of 0.0029 with a 95% confidence interval of 0.0017–0.0047 using mutations on chromosomes 16 and 21 (Arbeithuber *et al.* 2015). Direct measurement by estimation of these rates from *de novo* mutation studies in principle constitute a “gold standard”, subject to experimental limits and small sample sizes (Arbeithuber *et al.* 2015). Consistency of our population based estimates with these supports the robustness of our estimation procedure.

Our estimates for the slope in terms of the base pair mutation rate per centimorgan are somewhat greater than the overall figure of ∼ 1.5 × 10^−9^ obtained in Hellmann (2005). We note that Hellmann (2005) addressed the issue of spatial auto-correlation by means of the Cochrane-Orcutt correction (Kutner *et al.* 2005), which is applicable to an auto-regressive (AR) model. AR models are nested within ARMA models as a special case. Our analysis of the residuals (data not shown) demonstrates that, taking the case of chromosome 1, the optimal ARMA model has a superior AIC (∼ − 184, 554) to the optimal AR model (∼ − 184, 493). On the other hand, our estimate of the probability that a single recombination event causes a mutation is very different from the estimate in Hellmann (2005), which was also based on a population analysis. In contrast to our approach, which employs the intercept (see Results), this estimate was based on the slope. Specifically, the slope of the linear regression was estimated at ∼ 1.5 × 10^−9^ mutations per base pair per centimorgan, that is, every 1% increase in the recombination rate in 1 Mb of sequence generated an additional ∼ 0.0015 mutations per Mb. The number of mutation events per recombination event is then taken as 0.0015/0.01 = 0.15, which is two orders of magnitude greater than both the empirical estimates and our own. This argument implicitly treats all recombination-induced mutations as having been caused by recombination events in the most recent generation. It therefore ignores recombination-induced mutations caused by recombination events in previous generations.

The use of an ARMA distribution to model the residuals, rather than fitting a conventional linear regression, made a major difference to the results. To illustrate this, we repeated the analysis using an OLSLR model (Table S4). Comparing these estimates with those from the ARMA model (Table S1) shows marked differences in all parameter values. In particular, the variance due to recombination 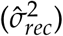 was some orders of magnitude larger in the OLSLR model. Critically, the intercept parameter was consistently lower for the OLSLR model and as a direct result the estimation of the number of mutations per recombination event was consistently greater for the OLSLR model by a factor of around 2. As correlations are given by the square root of *R*^2^ in a linear model, the discrepancies we identify raise doubts about the accuracy of estimates of correlation between recombination rates and substitution rates in studies that do not compensate for spatial auto-correlation (Lercher and Hurst 2002; Duret and Arndt 2008; Mugal and Ellegren 2011).

Our results indicate the effect of recombination depended on mutation direction with some mutations exhibiting no association (e.g. N→W transversions). This dependence on point mutation direction has been noted by other authors and our results are generally consistent with previous observations derived from substitution data (Duret and Arndt 2008). In particular, the mutation types seen to be most influenced by recombination were C→T and G→A, the same types that are subject to the CpG effect. In a study on *de novo* recombination and mutations (Arbeithuber *et al.* 2015), all but one of the 17 *de novo* mutations found in molecules with a crossover were of one of these two types (as were the three mutations found in non-crossover controls). A possible explanation is that recombination magnifies the CpG effect by effecting a temporary local strand separation, since the deamination of 5-methylcytosine is over 60 times more rapid in single-stranded than in double-stranded DNA (Ehrlich *et al.* 1986; Zhang and Mathews 1994).

Recombination will only contribute to variance in SNV density insofar as it occurs heterogeneously along the genome. Neither the proportion of mutation events occurring in a generation that are caused by recombination nor the probability that a recombination event gives rise to a mutation event (both ∼ 0.004) are affected by the distribution of recombination rates along the genome. For this reason, these quantities arguably provide a better measure of the direct effect of recombination on mutation. A comparative analysis of variance in SNV density due to recombination by chromosome given in Figure 1 showed some difference between chromosomes, with chromosomes 9, 15 and 17 having a significantly higher variance due to recombination than the other chromosomes. A possible explanation is that recombination rates are more heterogeneous along these chromosomes. One measure of heterogeneity is the variance of the recombination rates, which is shown in Figure 6. We see that chromosomes 15 and 17 do have relatively high variance, but not significantly higher than chromosome 13, while the variance for chromosome 21 is much greater. Chromosome 9, on the other hand, does not have a relatively high variance in recombination rate. Thus while variance in the recombination rate may contribute to variance in SNV density due to recombination, the explanation appears to be more complex. Another feature of Figure 1 is that chromosomes 9, 15 and 17 also have greater spread in their posterior distributions. This is likely to result directly from the fact that the variance due to recombination (*R*^2^) is greater in these cases (Wishart *et al.* 1931; Olkin and Finn 1995). In this and a number of other instances, chromosome 21 appears to be either an outlier or to take some extreme value. This may be an artefact due to the recombination map used having a much lower coverage of chromosome 21 (∼ 50%) than of other chromosomes (Kong *et al.* 2010, Supplementary Table 2).

**Figure 6.**
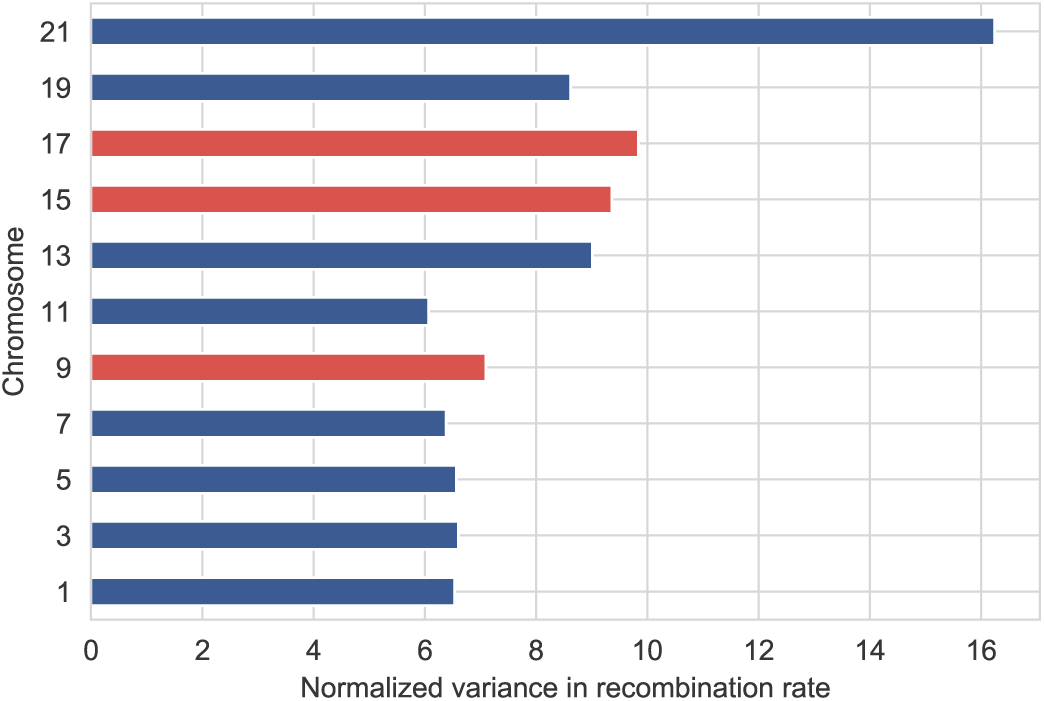
Variance in recombination rate by chromosome. The variance in recombination rate is calculated from average recombination rates for 10-kb bins, normalized by the mean recombination rate of the entire genome.

It has been argued that the relationship between recombination rate and SNV density is not causal. Some studies on *Drosophila*, while acknowledging the correlation between the density of SNVs and recombination, suggest that this is a result of the weaker effect of selective sweeps in regions of higher recombination, rather than a result of mutations being directly caused by crossover events (Begun and Aquadro 1992; McGaugh *et al.* 2012). The conclusion that recombination does not cause mutation has been used (Kern and Hahn 2018; Nachman 2001) to argue that the correlation between mutation and nucleotide diversity is proof of the near-ubiquitous occurrence of selective sweeps and hence of the action of selective forces throughout the genome, contrary to the tenets of the neutral theory (Kimura 1983). This interpretation is inconsistent with the empirical studies on *de novo* mutation and recombination (Arbeithuber *et al.* 2015). Additionally, our observation that the association between recombination and SNV density is dependent on mutation direction (Figure 2) has no apparent explanation in terms of selective sweeps.

### The influence of context on mutation

Our approach to analysing the influence of context on mutation has two main features: using a rich database of human variants identified by the 1KG Project (Aut 2015) and applying directly the concept of variance in mutation rate due to context as described in Materials and methods. This method is particularly suited to considering the issue of the effect on mutation of contexts of differing sizes.

A number of authors have variously dismissed (Hodgkinson *et al.* 2009; Krawczak *et al.* 1998) or made strong claims for (Aggarwala and Voight 2016; Zhu *et al.* 2017) the influence of contexts beyond 3-mers on mutation. Figure 3 shows that the increases from 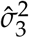 to 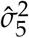 to 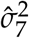 are relatively small. This is because the variance due to 3-mers incorporates the interaction between the central base of the 3-mer and its two flanking bases, which is the largest contributor to variance in SNV rates due to context. We found that ∼ 60% of 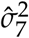 was due to the CpG effect.

For analysis of an individual point mutation, the central base is fixed and thus there are no explicit interactions with neighbouring bases. In this case, which we have referred to as fixed marginals (Figure 4), the additional variance added by 5-mers over 3-mers and, to an even greater extent, by 7-mers over 5-mers is substantial. These results may appear to support previous findings that the variance due to 7-mers is highly significant (Aggarwala and Voight 2016). However, the methods used in that work differ markedly to ours. When considering the influence of *k*-mers on the probability of observing an SNV, Aggarwala and Voight (2016) fitted a linear model to binomial data, which will not yield a valid maximum likelihood estimate of the slope and intercept parameters (Agresti 2002, p. 120). In calculating the relative influence of 3-mers and 7-mers, they used linear regression to predict the 7-mer SNV densities from the 3-mer SNV densities and calculated the *R*^2^ metric on this regression. This metric is mathematically the same as the ratio of the following quantities: sum of squares difference between the 3-mer SNV densities and the overall mean SNV density; and, the sum of squares difference between the 7-mer SNV densities and the overall mean SNV density. This ratio is in turn the ratio of variance in 3-mer SNV densities to that of 7-mer SNV densities (without being weighted by frequency of context.) This appears to account for some similarity of their results to ours.

In contrast to Aggarwala and Voight (2016), Zhu *et al.* (2017) used a log-linear model for estimating the information content of neighbouring bases. This approach is appropriate in modelling binomial data and also allowed comparison of the effect of different *k*-mers as measured by information content rather than traditional sum-of-squares variance. An advantage is that the joint effect of neighbouring nucleotides can be distinguished from the independent effects of each. That work likewise identified neighbouring nucleotides as distant as 4 bases away (hence *k* = 9) as associated with some transversion point mutations (Zhu *et al.* 2017).

The strong influence of 7-mer contexts apparent when conditioning on the central mutating base raises the question of whether this is due to specific hyper-mutable 7-mer contexts. Our investigation failed to identify any such 7-mer contexts that were not attributable to CpG hypermutability. The most mutagenic context was NNACGNN. This sequence is of course subject to the CpG effect and the 5’-A has a positive association on C→T mutations independent of the 3’ base (see Zhu *et al.* 2017, Figure 2). The incidence of ACG trinucleotides was ∼7% of that expected from the individual nucleotide frequencies.

Figure 5a provides evidence of strand-asymmetry in the variance due to contextual influence for all 12 point mutation directions. It has been conjectured that such strand-asymmetry is caused by transcription coupled DNA repair (TCR) (e.g. Hwang and Green 2004). TCR is a strand-asymmetric process which occurs in actively transcribed genes when an RNA polymerase (RNAP) translocating along a DNA strand encounters a distorting lesion or other local factor that retards its forward progress and may cause it to recruit nucleotide excision repair proteins (Spivak and Ganesan 2014). Sequence context is known to be involved in factors that can pause or arrest RNAPs (Spivak and Ganesan 2014). In their phylogenetic analysis of substitution rates, Hwang and Green (2004) found that T→C substitution rates were higher than those for A→G and C→T substitution rates were higher than those for G→A. Our analysis of SNV data differs from this in showing A→G to have a significantly higher SNV density than T→C while C→T SNVs had only marginally greater SNV density compared to G→A (Table S3). Figure 5a shows the same pattern in variance due to context: 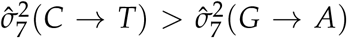 and 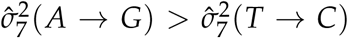. Hwang and Green (2004) also showed transcription-associated mutational asymmetry to be influenced by context for transitions. Our results indicate that such influence occurs to some significant degree for all mutation directions. Overall, it appears that a substantial association exists between TCR and variance in SNV density.

## Conclusion

We have demonstrated that estimating the variance in SNV density due to context can discriminate the effect of contexts of different sizes. This was done from three perspectives: considering the 12 point mutation directions separately; aggregating over these directions while marginalizing over the central allele; and aggregating over these directions without marginalizing over the central allele (measured by 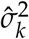). The perspective adopted has a marked influence on estimates of relative influence. For example, results aggregated over mutation direction will be dominated by the more abundant transition mutations and in particular, by the CpG effect. Our approach has clarified the relationship between results from these different perspectives and, in particular, has demonstrated the dominant effect of the interaction between a central allele and its immediate neighbours. The use of Bayesian posterior distributions was able to give a high degree of certainty to conclusions about the strand-asymmetry of contextual influence in intronic regions. Further, our methods are driven solely by varying SNV densities between contexts and are not influenced by the distribution of *k*-mers within the genome.

We also quantified variance in SNV density due to recombination. However, a direct comparison of this quantity with the variance in SNV density due to context has some limitations. We measured variance in SNV density due to recombination at the scale of 10-kb DNA blocks. This does not take account of any variance due to recombination that exists within 10-kb blocks. This limitation is not easily overcome as there is presently no data for fine scale recombination at the individual base level. We note that the quantitative impacts of recombination and context on mutation are conceptually difficult to compare meaningfully, as context is a state and recombination an event. For this reason, the proportion of mutation events caused by recombination and the probability that a recombination event gives rise to a mutation event (both ∼ 0.004) are better measures of the direct impact of recombination on mutation. Our estimate that recombination only accounts for ∼ 0.4% of the average mutation rate makes recombination appear a relatively minor contributor to mutation rate overall. However, recombination is concentrated in hotspots, typically 1 - 2 kb in length, in which the recombination rate can commonly be 50 or more times higher than average (Altshuler and Donnelly 2005). In such regions, recombination would account for ∼ 20% or more of the mutation rate.

## Acknowledgment

This research is supported by an Australian Government Research Training Program (RTP) Scholarship to HS.

## Supplementary for “Sequence context, recombination rate and the probability of observing point mutations”

**Table S1:** Results of analysis of variance due to recombination by chromosome for odd-numbered chromosomes. ‘p’ and ‘q’ define the ARMA(p,q) distribution used; ‘Slope’ and ‘Intercept’ are the estimated parameters of the linear model expressed in terms of change in SNV density per centimorgan and SNV density respectively; 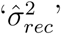 is the estimated variance in SNV density due to recombination; ‘Slope (M)’ is the estimated slope parameter expressed as change in mutation rate per centimorgan; and ‘Mutations’ is the estimated average number of mutations resulting from a recombination event.

| Chr | SNV density | p | q | Slope | Intercept | $\hat{\sigma}_{rec}^2$ | Slope (M) | Mutations |
| --- | --- | --- | --- | --- | --- | --- | --- | --- |
| 1 | 0.0252 | 8 | 2 | 0.0085 | 0.0251 | 3.08e-07 | 3.67e-09 | 0.0041 |
| 3 | 0.0262 | 9 | 4 | 0.0072 | 0.0261 | 1.87e-07 | 3.39e-09 | 0.0038 |
| 5 | 0.0261 | 8 | 4 | 0.0075 | 0.0260 | 1.76e-07 | 3.42e-09 | 0.0039 |
| 7 | 0.0262 | 9 | 2 | 0.0088 | 0.0261 | 3.07e-07 | 4.03e-09 | 0.0049 |
| 9 | 0.0254 | 4 | 4 | 0.0085 | 0.0253 | 6.26e-07 | 3.51e-09 | 0.0043 |
| 11 | 0.0269 | 3 | 3 | 0.0087 | 0.0268 | 1.57e-07 | 3.88e-09 | 0.0042 |
| 13 | 0.0261 | 7 | 4 | 0.0080 | 0.0260 | 2.22e-07 | 3.06e-09 | 0.0031 |
| 15 | 0.0261 | 2 | 2 | 0.0079 | 0.0260 | 4.88e-07 | 3.13e-09 | 0.0032 |
| 17 | 0.0256 | 10 | 1 | 0.0091 | 0.0255 | 6.02e-07 | 4.06e-09 | 0.0039 |
| 19 | 0.0275 | 8 | 4 | 0.0071 | 0.0274 | 1.91e-07 | 3.08e-09 | 0.0033 |
| 21 | 0.0275 | 2 | 2 | 0.0067 | 0.0274 | 1.73e-07 | 2.13e-09 | 0.0022 |

**Table S2:** Posterior probability that recombination does not have a positive effect on mutation by point mutation direction and chromosome (odd-numbered chromosomes shown).

|  | 1 | 3 | 5 | 7 | 9 | 11 | 13 | 15 | 17 | 19 | 21 |
| --- | --- | --- | --- | --- | --- | --- | --- | --- | --- | --- | --- |
| C→T | 0.00 | 0.00 | 0.00 | 0.00 | 0.00 | 0.00 | 0.00 | 0.00 | 0.00 | 0.00 | 0.00 |
| G→A | 0.00 | 0.00 | 0.00 | 0.00 | 0.00 | 0.00 | 0.00 | 0.00 | 0.00 | 0.00 | 0.03 |
| T→C | 0.00 | 0.00 | 0.00 | 0.00 | 0.00 | 0.00 | 0.00 | 0.00 | 0.00 | 0.01 | 0.00 |
| A→G | 0.00 | 0.00 | 0.00 | 0.00 | 0.00 | 0.00 | 0.00 | 0.00 | 0.00 | 0.00 | 0.00 |
| C→G | 0.00 | 0.00 | 0.00 | 0.32 | 0.03 | 0.41 | 0.00 | 0.66 | 0.02 | 0.06 | 0.54 |
| G→C | 0.00 | 0.20 | 0.00 | 0.00 | 0.01 | 0.01 | 0.03 | 0.52 | 0.05 | 0.46 | 0.00 |
| T→G | 0.04 | 0.01 | 0.00 | 0.03 | 0.34 | 0.53 | 0.00 | 0.15 | 0.12 | 0.81 | 0.18 |
| A→C | 0.01 | 0.00 | 0.00 | 0.00 | 0.04 | 0.45 | 0.01 | 0.20 | 0.00 | 0.07 | 0.26 |
| T→A | 0.71 | 0.77 | 0.28 | 0.87 | 0.38 | 0.42 | 0.09 | 0.52 | 0.21 | 0.47 | 0.89 |
| A→T | 0.50 | 0.99 | 0.53 | 0.63 | 0.22 | 0.31 | 0.98 | 0.04 | 0.09 | 0.00 | 0.00 |
| C→A | 0.97 | 1.00 | 0.99 | 0.70 | 0.37 | 0.90 | 0.34 | 0.09 | 0.07 | 0.85 | 0.01 |
| G→T | 0.95 | 0.65 | 0.95 | 0.78 | 0.99 | 0.91 | 0.79 | 0.13 | 0.86 | 0.95 | 0.79 |

**Table S3:** Variance in probability of SNVs due to context. 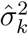 denote the estimated variances for context size *k*. The size of context includes the central allele. Results are conditioned on mutation direction (ancestral and derived state). The column ‘Density’ shows the density for each SNV direction (conditioned on the ancestral allele) for reference. See Methods and materials for data sources.

| **Mutation** | **Density** | $\hat{\sigma}_3^2$ | $\hat{\sigma}_5^2$ | $\hat{\sigma}_7^2$ |
| --- | --- | --- | --- | --- |
| C→T | 0.0208 | 6.20e-04 | 6.75e-04 | 7.99e-04 |
| C→A | 0.0044 | 5.95e-06 | 9.27e-06 | 1.37e-05 |
| C→G | 0.0052 | 1.07e-05 | 1.48e-05 | 2.11e-05 |
| T→C | 0.0093 | 8.33e-06 | 2.09e-05 | 2.96e-05 |
| T→A | 0.0024 | 4.96e-07 | 1.53e-06 | 3.21e-06 |
| T→G | 0.0029 | 6.29e-07 | 1.18e-06 | 2.37e-06 |
| A→C | 0.0029 | 3.63e-07 | 7.68e-07 | 1.57e-06 |
| A→T | 0.0028 | 6.81e-07 | 1.69e-06 | 3.43e-06 |
| A→G | 0.0135 | 2.03e-05 | 6.04e-05 | 7.48e-05 |
| G→C | 0.0044 | 6.91e-06 | 1.01e-05 | 1.45e-05 |
| G→T | 0.0050 | 8.08e-06 | 1.24e-05 | 1.83e-05 |
| G→A | 0.0203 | 5.13e-04 | 5.58e-04 | 6.65e-04 |

**Figure S1:**
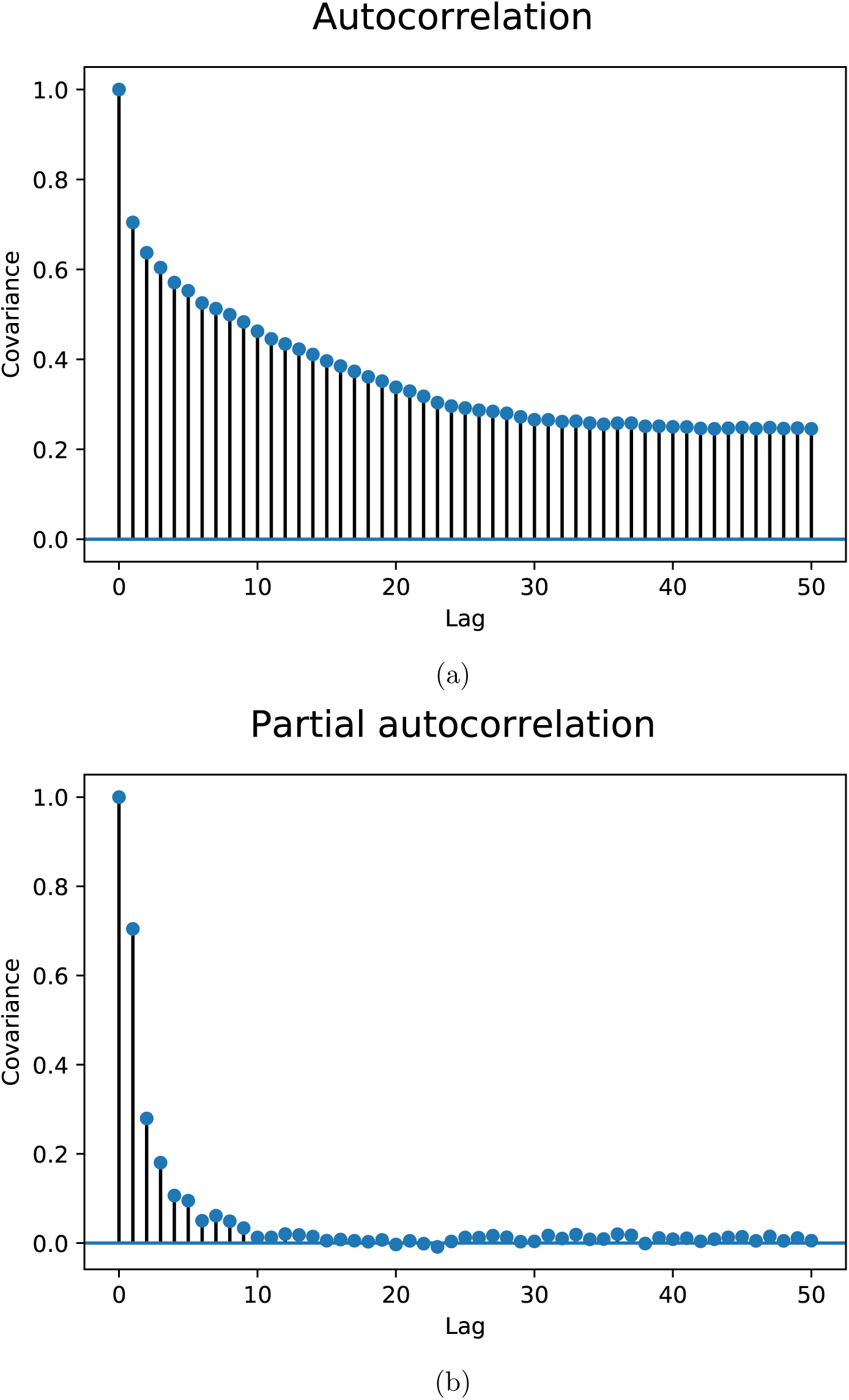
Visualisation of auto-correlation of residuals from ordinary least squares linear regression of SNV densities against average recombination rates for chromosome 1. Correlation between residuals in bins separated by lags in the range 0 to 50 from (a) auto-correlation. (b) partial auto-correlation. The analysis removed the effect of correlations at shorter lags and indicates the number of lags required in an auto-regressive model (10 in this instance). The blue shading shows a 95% significance interval.

**Figure S2:**
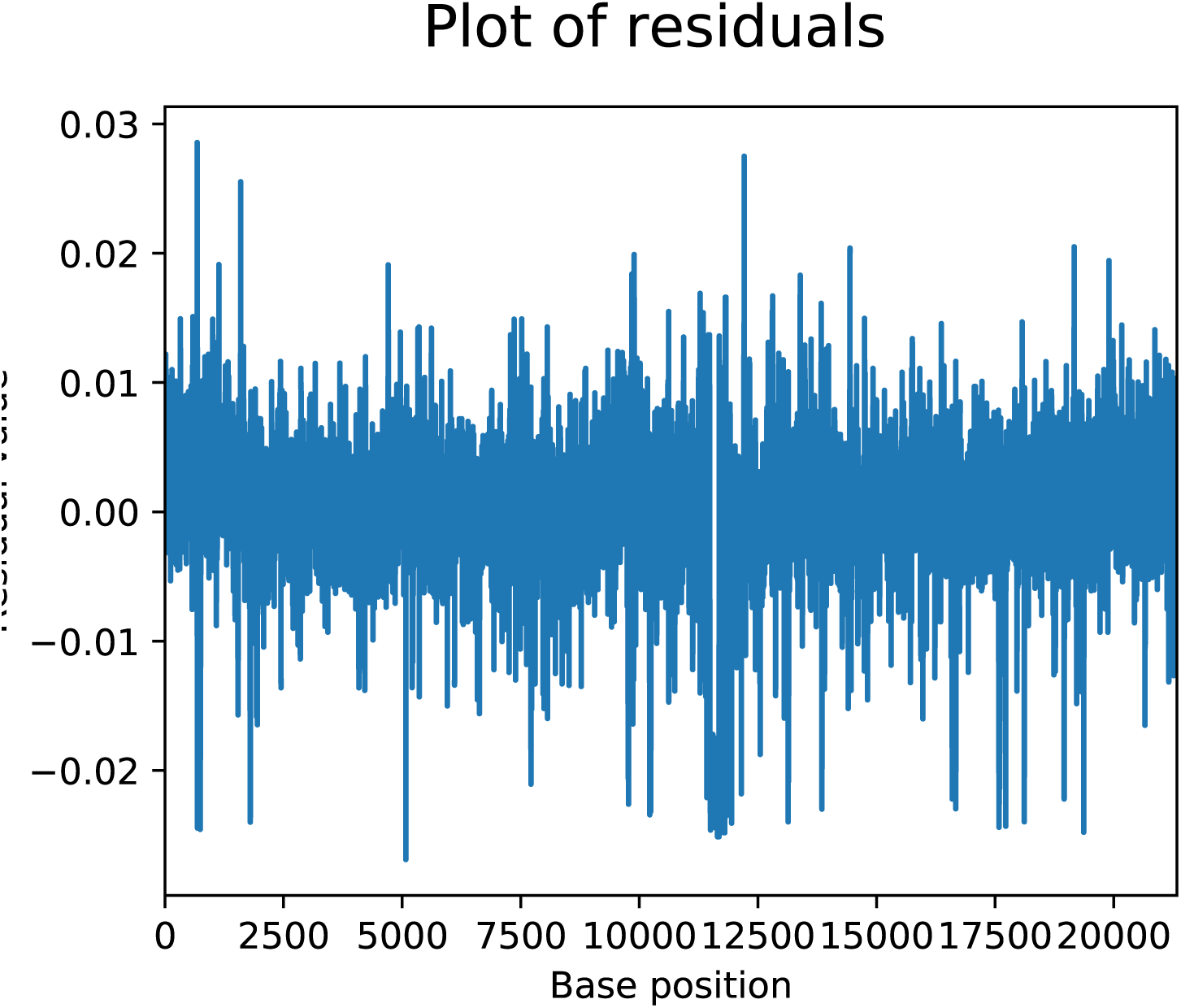
Visualisation of residuals of ordinary least squares linear regression of SNV densities against average recombination rates for chromosome 1 for 10-kb bins spanning the length of the chromosome. The residuals appear to have a constant mean of zero and a constant variance as required by stationarity. Stationarity is formally demonstrated by the Dickey-Fuller test.

**Table S4:** Results of analysis of variance due to recombination by chromosome for odd-numbered chromosomes using ordinary last squares linear regression (OLSLR). ‘Slope’ and ‘Intercept’ are the estimated parameters of the linear model expressed in terms of change in SNV density per centimorgan and SNV density respectively; 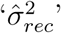 is the estimated variance in SNV density due to recombination; and ‘Mutations’ is the estimated average number of mutations resulting from a recombination event.

| Chr | SNV density | Slope | Intercept | $\hat{\sigma}_{rec}^2$ | Mutations |
| --- | --- | --- | --- | --- | --- |
| 1 | 0.0252 | 0.0003 | 0.0249 | 0.0225 | 0.0107 |
| 3 | 0.0262 | 0.0002 | 0.0260 | 0.0193 | 0.0082 |
| 5 | 0.0261 | 0.0002 | 0.0259 | 0.0123 | 0.0075 |
| 7 | 0.0262 | 0.0003 | 0.0259 | 0.0187 | 0.0111 |
| 9 | 0.0254 | 0.0005 | 0.0249 | 0.0262 | 0.0179 |
| 11 | 0.0269 | 0.0002 | 0.0267 | 0.0096 | 0.0067 |
| 13 | 0.0261 | 0.0002 | 0.0259 | 0.0245 | 0.0060 |
| 15 | 0.0261 | 0.0003 | 0.0257 | 0.0315 | 0.0124 |
| 17 | 0.0256 | 0.0003 | 0.0252 | 0.0454 | 0.0150 |
| 19 | 0.0275 | 0.0002 | 0.0272 | 0.0093 | 0.0076 |
| 21 | 0.0275 | 0.0001 | 0.0274 | 0.0172 | 0.0036 |

## Literature Cited

2015 A global reference for human genetic variation. Nature 526: 68–74.

Aggarwala, V. and B. Voight, 2016 An expanded sequence context model broadly explains variability in polymorphism levels across the human genome. Nat Genet 48: 349–55.

Agresti, A., 2002 Categorical Data Analysis. John Wiley & Sons, second edition.

Altshuler, D. and P. Donnelly, 2005 A haplotype map of the human genome. Nature 437: 1299.

Arbeithuber, B., A. J. Betancourt, T. Ebner, and T. Irene, 2015 Crossovers are associated with mutation and biased gene conversion at recombination hotspots. Proc National Acad Sci 112: 2109–2114.

Bayer, M., 2012 Sqlalchemy. In The Architecture of Open Source Applications Volume II: Structure, Scale, and a Few More Fearless Hacks, edited by A. Brown and G. Wilson, aosabook.org.

Begun, D. J. and C. F. Aquadro, 1992 Levels of naturally occurring DNA polymorphism correlate with recombination rates in D. melanogaster. Nature 356: 519.

Benzer, S., 1961 On the topography of the genetic fine structure. Proceedings of the National academy of Sciences of the United States of America 47: 403.

Bulmer, M., 1986 Neighboring base effects on substitution rates in pseudogenes. Molecular biology and evolution 3: 322–329.

Coulondre, C., J. H. Miller, P. J. Farabaugh, and W. Gilbert, 1978 Molecular basis of base substitution hotspots in Escherichia coli. Nature 274: 775–780.

Cunningham, F., M. Amode, D. Barrell, K. Beal, K. Billis, et al., 2015 Ensembl 2015. Nucleic Acids Res 43: D662–9.

Cuny, G., P. Soriano, G. Macaya, and G. Bernardi, 1981 The major components of the mouse and human genomes: 1. Preparation, basic properties and compositional heterogeneity. European Journal of Biochemistry 115: 227–233.

Duret, L. and P. F. Arndt, 2008 The impact of recombination on nucleotide substitutions in the human genome. PLoS Genetics 4: e1000071.

Ehrlich, M., K. F. Norris, R. Y. Wang, K. C. Kuo, and C. W. Gehrke, 1986 DNA cytosine methylation and heat-induced deamination. Bioscience reports 6: 387–393.

Ehrlich, M. and R. Wang, 1981 5-Methylcytosine in eukaryotic DNA. Science 212: 1350–1357.

Gojobori, T., W.-H. Li, and D. Graur, 1982 Patterns of nucleotide substitution in pseudogenes and functional genes. Journal of Molecular Evolution 18: 360–369.

Hellmann, I., 2005 Why do human diversity levels vary at a megabase scale? Genome Research 15: 1222–1231.

Hodgkinson, A. and A. Eyre-Walker, 2011 Variation in the mutation rate across mammalian genomes. Nature Reviews Genetics 12: 756–766.

Hodgkinson, A., E. Ladoukakis, and A. Eyre-Walker, 2009 Cryptic variation in the human mutation rate. PLoS Biol 7: e1000027.

Hunter, J. D., 2007 Matplotlib: A 2D graphics environment. Computing in Science & Engineering 9: 90–95.

Huttley, G., 2016 scitrack 0.1.3. https://pypi.org/project/scitrack/0.1.3.

Huttley, G. and H. Ying, 2009 ensembldb3. https://github.com/cogent3/ensembldb3.

Hwang, D. G. and P. Green, 2004 Bayesian Markov chain Monte Carlo sequence analysis reveals varying neutral substitution patterns in mammalian evolution. Proceedings of the National Academy of Sciences 101: 13994–14001.

Jónsson, H., P. Sulem, B. Kehr, S. Kristmundsdottir, F. Zink, et al., 2017 Parental influence on human germline de novo mutations in 1,548 trios from Iceland 549: 519–522.

Kern, A. D. and M. W. Hahn, 2018 The neutral theory in light of natural selection. Molecular biology and evolution 35: 1366–1371.

Kimura, M., 1983 The neutral theory of molecular evolution. Cambridge University Press.

Knight, R., P. Maxwell, A. Birmingham, J. Carnes, J. Caporaso, et al., 2007 PyCogent: a toolkit for making sense from sequence. Genome Biol 8: R171.

Kong, A., G. Thorleifsson, D. F. Gudbjartsson, G. Masson, A. Sigurdsson, et al., 2010 Fine-scale recombination rate differences between sexes populations and individuals. Nature 467: 1099–1103.

Krawczak, M., E. V. Ball, and D. N. Cooper, 1998 Neighboring-nucleotide effects on the rates of germ-line single-base-pair substitution in human genes. The American Journal of Human Genetics 63: 474–488.

Kutner, M. H., C. J. Nachtsheim, J. Neter, W. Li, et al., 2005 Applied linear statistical models. McGraw-Hill New York.

Lercher, M. J. and L. D. Hurst, 2002 Human SNP variability and mutation rate are higher in regions of high recombination. Trends in Genetics 18: 337–340.

McGaugh, S. E., C. S. Heil, B. Manzano-Winkler, L. Loewe, S. Goldstein, et al., 2012 Recombination modulates how selection affects linked sites in Drosophila. PLoS biology 10: e1001422.

McKinney, W., 2010 Data structures for statistical computing in Python. In Proceedings of the 9th Python in Science Conference, edited by S. van der Walt and J. Millman, pp. 51–56.

Mills, T. C., 2008 The Econometric Modelling of Financial Time Series. Cambridge University Press, third edition.

Mizon, G. E., 1995 A simple message for autocorrelation correctors: Don’t. Journal of Econometrics 69: 267–288.

Molenberghs, G., G. Fitzmaurice, M. G. Kenward, A. Tsiatis, and G. Verbeke, 2014 Handbook of missing data methodology. Chapman and Hall/CRC.

Mugal, C. F. and H. Ellegren, 2011 Substitution rate variation at human CpG sites correlates with non-CpG divergence, methylation level and GC content. Genome biology 12: R58.

Nachman, M. W., 2001 Single nucleotide polymorphisms and recombination rate in humans. TRENDS in Genetics 17: 481–485.

Olkin, I. and J. D. Finn, 1995 Correlations redux. Psychological Bulletin 118: 155.

Pedregosa, F., G. Varoquaux, A. Gramfort, V. Michel, B. Thirion, et al., 2011 Scikit-learn: Machine learning in Python. Journal of Machine Learning Research 12: 2825–2830.

Ramsahoye, B. H., D. Biniszkiewicz, F. Lyko, V. Clark, A. P. Bird, et al., 2000 Non-CpG methylation is prevalent in embryonic stem cells and may be mediated by DNA methyltransferase 3a. Proceedings of the National Academy of Sciences 97: 5237–5242.

Roach, J. C., G. Glusman, A. F. A. Smit, C. D. Huff, R. Hubley, et al., 2010 Analysis of genetic inheritance in a family quartet by whole-genome sequencing. Science 328: 636–639.

Ronacher, A., 2009 click 7.0. https://pypi.org/project/click/.

Salvatier, J., T. V. Wiecki, and C. Fonnesbeck, 2016 Probabilistic programming in Python using PyMC3. PeerJ Computer Science 2: e55.

Seabold, S. and J. Perktold, 2010 statsmodels: Econometric and statistical modeling with Python. In 9th Python in Science Conference.

Smith, T. C., P. F. Arndt, and A. Eyre-Walker, 2018 Large scale variation in the rate of germ-line de novo mutation, base composition, divergence and diversity in humans. PLoS genetics 14: e1007254.

Spivak, G. and A. K. Ganesan, 2014 The complex choreography of transcription-coupled repair. DNA repair 19: 64–70.

Theano Development Team, 2016 Theano: A Python framework for fast computation of mathematical expressions. arXiv e-prints abs/1605.02688.

Tretyakov, K., 2013 pyliftover 0.4. https://pypi.org/project/pyliftover/.

Virtanen, P., R. Gommers, T. E. Oliphant, M. Haberland, T. Reddy, et al., 2019 SciPy 1.0–Fundamental Algorithms for Scientific Computing in Python. arXiv e-prints p. 1907.10121.

Waskom, M., O. Botvinnik, D. O’Kane, P. Hobson, S. Lukauskas, et al., 2017 Seaborn: v0.8.1. https://doi.org/10.5281/zenodo.883859.

Wishart, J., T. Kondo, and E. Elderton, 1931 The mean and second moment coefficient of the multiple correlation coefficient, in samples from a normal population. Biometrika pp. 353–376.

Ying, H., J. Epps, R. Williams, and G. Huttley, 2010 Evidence that localized variation in primate sequence divergence arises from an influence of nucleosome placement on DNA repair. Molecular biology and evolution 27: 637–649.

Zhang, X. and C. Mathews, 1994 Effect of DNA cytosine methylation upon deamination-induced mutagenesis in a natural target sequence in duplex DNA. Journal of Biological Chemistry 269: 7066–7069.

Zhu, Y., T. Neeman, V. B. Yap, and G. A. Huttley, 2017 Statistical methods for identifying sequence motifs affecting point mutations. Genetics 205: 843–856.

